# Individual Stability of Pain- and Touch-Related Neuronal Gamma Oscillations

**DOI:** 10.1101/2021.11.21.469419

**Authors:** Elia Valentini, Alina Shindy, Viktor Witkovsky, Anne Stankewitz, Enrico Schulz

## Abstract

**Background:** The processing of brief pain and touch stimuli has been associated with an increase of neuronal oscillations in the gamma range (40-90 Hz). However, some studies report divergent gamma effects across single participants.

**Methods:** In two repeated sessions we recorded gamma responses to pain and touch stimuli using EEG. Individual gamma responses were extracted from EEG channels and from ICA components that contain a strong gamma amplitude.

**Results:** We observed gamma responses in the majority of the participants. If present, gamma synchronisation was always bound to a component that contained a laser-evoked response. We found a broad variety of individual cortical processing: some participants showed a clear gamma effect, others did not exhibit any gamma. For both modalities, the effect was reproducible between sessions. In addition, participants with a strong gamma response showed a similar time-frequency pattern across sessions.

**Conclusions:** Our results indicate that current measures of reproducibility of research results do not reflect the complex reality of the diverse individual processing pattern of applied pain and touch. The present findings raise the question of whether we would find similar quantitatively different processing patterns in other domains in neuroscience: group results would be replicable but the overall effect is driven by a subgroup of the participants.

## Introduction

Research spanning several decades has highlighted the crucial role of high frequency brain oscillations in perceptual integration [Pritchett et al., 2015]. A number of these publications focused on the investigation of pain-related and touch-related neuronal oscillations in the gamma range [Heid et al., 2020; Michail et al., 2016; Schulz et al., 2015; Valentini et al., 2013]. These studies showed an increase of gamma activity in response to laser pain at around 80 Hz at central and lateral electrodes [Hauck et al., 2015; Schulz et al., 2012] and from intracerebral recordings from the insular cortex [Hauck et al., 2017; Liberati et al., 2018].

There are fewer studies on gamma responses to brief touch [Michail et al., 2016] and more extended vibrotactile stimulation [Ryun et al., 2017]. Gamma oscillations, however, were found to encode the intensity of brief phasic pain stimulation [Hauck et al., 2007; Schulz et al., 2011] and long-lasting, tonic pain [May et al., 2018; Nickel et al., 2017; Schulz et al., 2015]. Some publications sparked controversial responses claiming that gamma oscillations are the only neuronal oscillation predictive of the perception of pain [Zhang et al., 2012]. However, others recently questioned the neuronal origin of gamma oscillations in response to noxious stimuli [Chouchou et al., 2021]. There are two potential reasons that fuel this controversy:

### Insufficient artefact cleaning

Firstly, EEG responses to painful laser application in the gamma range are prone to event-related muscle artefacts [Muthukumaraswamy, 2010; Yuval-Greenberg et al., 2008]. These artefacts can obscure any potential effect of cortical origin. Therefore, a thorough cleaning with a single trial inspection of the time-frequency transformed data (preferably on data decomposed with independent component analysis, ICA) is of utmost importance [Liberati et al., 2018; Michail et al., 2016]. Consequently, studies that do not sufficiently do so will inevitably exhibit effects that resemble the topography of neck or face muscles [Chouchou et al., 2021]. The use of a multi-electrode array is imperative in order to assess the plausibility of the effect by evaluating gamma topographies [Muthukumaraswamy, 2010]. Everything below these standards can raise legitimate doubts on the cortical origin of the gamma response [Yuval-Greenberg et al., 2008]. This issue can be solved with appropriate data processing [Liebisch et al., 2021; Nottage et al., 2013] and logical reasoning [Fries et al., 2008; Muthukumaraswamy, 2010].

### Interindividual variable response patterns and absence of gamma oscillations

Secondly, some researchers may have failed to detect gamma oscillations in response to painful laser application. Although gamma effects at group level have been repeatedly published [Hauck et al., 2017; Schulz et al., 2011], individual data show that not every participant has a clear effect at about 80 Hz. Indeed, the number of subjects per study that exhibit gamma oscillations can vary substantially. For one study, most participants show event-related gamma oscillations in response to pain and touch stimuli [Michail et al., 2016]. In stark contrast, in a sample of fibromyalgia patients and age-matched controls [Tiemann et al., 2012], none of the 40 participants showed any gamma response (unpublished aspects of the study series). This variability also applies to studies on long-lasting pain [May et al., 2018; Schulz et al., 2015], which reported a positive relationship between the magnitude of gamma and pain intensity; they confirm the variability of individual pain-related gamma oscillations by also showing no relationship or even a negative relationship between gamma activity and pain intensity for some subjects at frontocentral electrodes [May et al., 2018; Schulz et al., 2015].

While the first point highlights the responsibility and skills of the researcher, we can address the second point and aim to investigate in greater depth the broad range of individual pain- and touch-related responses in the gamma range. However, previous studies did not address this variation of gamma event-related oscillations and the individual stability across subjects in repeated recordings. We hypothesised that participants with a clear gamma response exhibit this response in either session of pain and touch stimulation. Subjects with a low gamma amplitude should exhibit this effect in both sessions. Furthermore, we hypothesised no overall change of the participants’ gamma amplitudes across both sessions.

## Methods

### Subjects

The raw data used in the current work were previously reported in two distinct studies [Michail et al., 2016; Tiemann et al., 2014]. EEG was recorded from 22 healthy male human subjects with a mean age of 24 years (21–31 years). All subjects gave written informed consent. The study was approved by the local ethics committee and conducted in conformity with the Declaration of Helsinki.

### Paradigm

In two counterbalanced sessions, a total of 150 painful cutaneous laser stimuli and 150 touch stimuli of matched intensities were delivered to the dorsum of the right hand (Figure 1). The laser device used was a Tm:YAG laser (Starmedtec GmbH, Starnberg, Germany) with a wavelength of 1960 nm, a pulse duration of 1 ms and a spot diameter of 5 mm. The physical energy of the pain stimulation was kept constant at 600 mJ. To prevent skin damage, the stimulation site was changed slightly after each stimulus. Touch stimuli with a force of 181 mN were applied using von Frey monofilaments delivered through a computer-controlled device as described in detail previously [Dresel et al., 2008; Michail et al., 2016].

**Figure 1.**
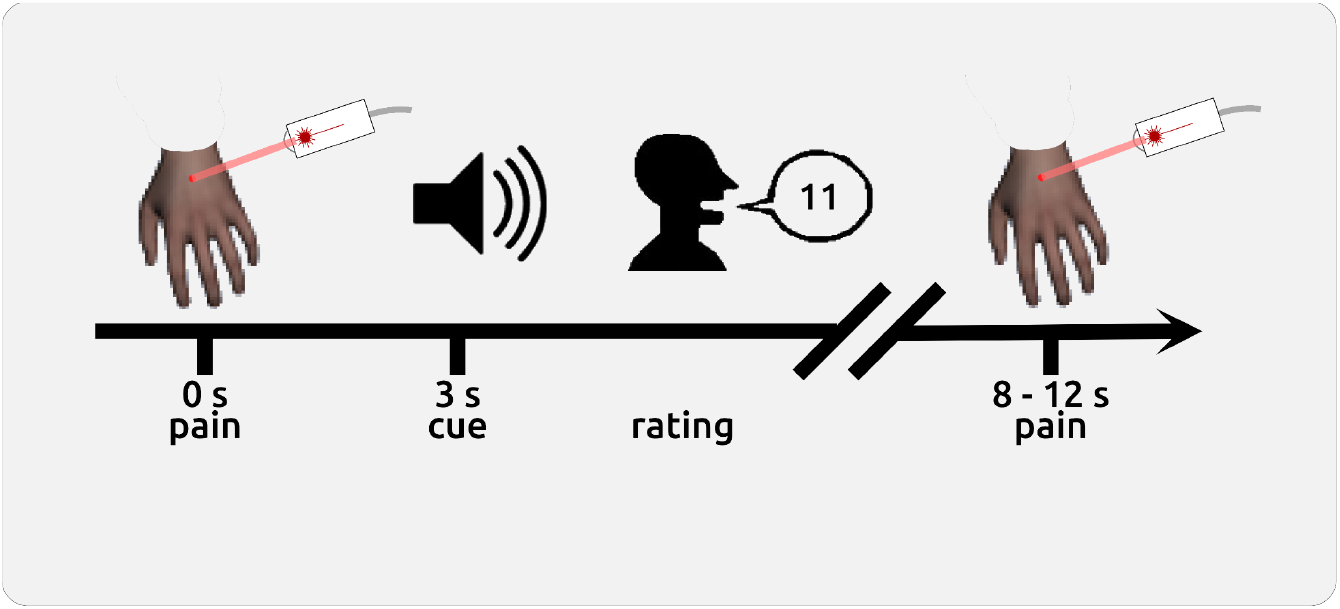
Paradigm and behavioural results. 75 laser stimuli were applied with an interstimulus interval of 8 to 12 s. Three seconds after each stimulus, an auditory cue prompted the subject to rate the pain intensity. The same procedure was utilised for touch stimuli.

Interstimulus intervals (ISI) for both modalities were randomly varied between 8 and 12 seconds. The subjects perceived the stimuli with closed eyes. Three seconds after each stimulus, the subjects were prompted by an auditory cue to verbally rate the perceived intensity of the stimulus on a 0-10 numerical rating scale. For pain stimuli, this was anchored by no pain (0) and maximum pain (10) the subjects were willing to tolerate during the experiment. For touch stimuli, the scale ranged between no perception (0) and maximal imaginable touch (10) that was not perceived as painful. The entire procedure of dopamine depletion, which neither significantly affected the perception of pain intensity nor the magnitude of gamma oscillations, has been described elsewhere [Tiemann et al., 2014]. The re-analysis of the current data involves a new integrated statistical modelling as well as advanced handling of muscle artefacts interfering with the genuine interpretation of the neural origins of the gamma oscillations [Liebisch et al., 2021].

### EEG recording and preprocessing

EEG data were recorded using an MRI-compatible electrode cap (FMS, Munich, Germany). The electrode montage included 64 electrodes, was referenced to the FCz electrode, grounded at AFz, and sampled at 1 kHz with 0.1 μV resolution. Impedance was kept below 20 kΩ. Raw subject-concatenated EEG data were preprocessed in Brain Vision Analyzer software (Brain Products, Munich, Germany). After the application of a 0.5 Hz high-pass filter, data were decomposed into 64 components using ICA. Components related to horizontal and vertical eye movements, as well as major artefacts, were removed and data were back-transformed into EEG signals. Muscle artefacts were removed using a dedicated algorithm [Liebisch et al., 2021]. Epoched artefact-free data of each EEG electrode were downsampled to 512 Hz, re-referenced to the common average reference, and time-frequency decomposed (for details see [Michail et al., 2016]. Single-trial z-transformed EEG data were finally checked for remaining high-frequency artefacts.

### Statistical Analysis

#### Electrode-based analysis

Based on our previous findings; we defined the gamma windows for the analysis on EEG electrodes between 76-86 Hz and 0.15-0.35s [Michail et al., 2016; Schulz et al., 2012]. Gamma responses were computed as percent signal change in reference to the prestimulus baseline (−1000 to −100 ms) for each trial. In EEG space, we utilised a conventional approach and conducted statistical analysis separately for each electrode. However, this approach can not take individual topographical variability of the gamma responses into account as it relies on information collected by single sensors. To take into account the whole scalp topographical information, we spatially filtered the data using ICA across all pain and touch trials.

#### Independent component-based analysis

We first conducted a time-frequency (TF) analysis on epoched ICA components using the same parameters as for the EEG channels. Most participants exhibited one to four components with a clear gamma response. All components with gamma responses were related to a component with evoked activity, had an interpretable topography, and were artefact-free. For each subject, we selected the component with the strongest gamma response. This approach has two advantages. *First*, we take individual variation into account. The present data show a tremendous variability of gamma responses with distinct topographical distributions and unique but replicable TF ranges. *Second*, these components are completely artefact free and would not need any additional treatment (see Supplementary 1).

Individual gamma time-frequency windows were defined from averaged single-trial oscillatory responses and included data points with at least 15% signal change (compared to the prestimulus baseline) in the gamma range in the averaged data across all 300 trials. Based on these super-threshold averaged data points, single-trial gamma data were computed. For subjects without gamma components, we extracted the single-trial gamma magnitude from the component with the largest evoked response within the literature-based TF window (76-86Hz; 150-350ms).

#### Statistical tests on stability, habituation, and order

We separately ran the statistical analyses for both pain and touch. All tests were performed on three dependent variables: (a) *behavioural* data, (b) gamma responses from *EEG* electrodes, and (c) gamma responses from *ICA* components.

*First*, the combined consideration of the between-subject correlation and the potential effect of order provides a measure of stability. To this aim, we computed Kendall’s τ coefficient on averaged rating and gamma data to test whether the subjects with high parameter values in the first session exhibit high parameter values in the second session and vice versa:

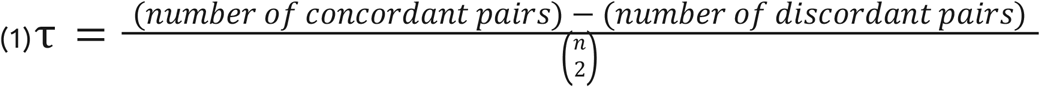

*Second*, using linear mixed effect models (LMEs), we tested whether there is a dependency of single-trial stimulus ratings or single-trial gamma responses on the order of testing sessions or the depletion intervention:

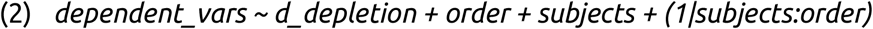

*Third*, using LMEs, we tested whether our dependent variables (ratings, ICA-gamma, EEG-gamma) habituate over time. The vector to represent habituation was defined according to the trial number per session (from 1 to 75).

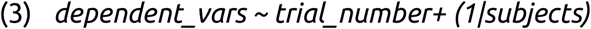

#### Multiple testing

To correct for multiple testing for the analyses on EEG data, we applied a randomisation approach. Depending on the analysis, rating data or the coding of order and depletion were shuffled (function “randperm” in Matlab) and the entire analysis was repeated 5000 times. The highest absolute t-values of each repetition across all electrodes were extracted. This procedure resulted in a right-skewed distribution of the 5000 max values for each analysis. Based on these distributions, the statistical thresholds were determined using the “palm_datapval” function publicly available in PALM [Winkler et al., 2014].

## Results

Table 1 provides a summary of the statistical findings.

**Table 1.**
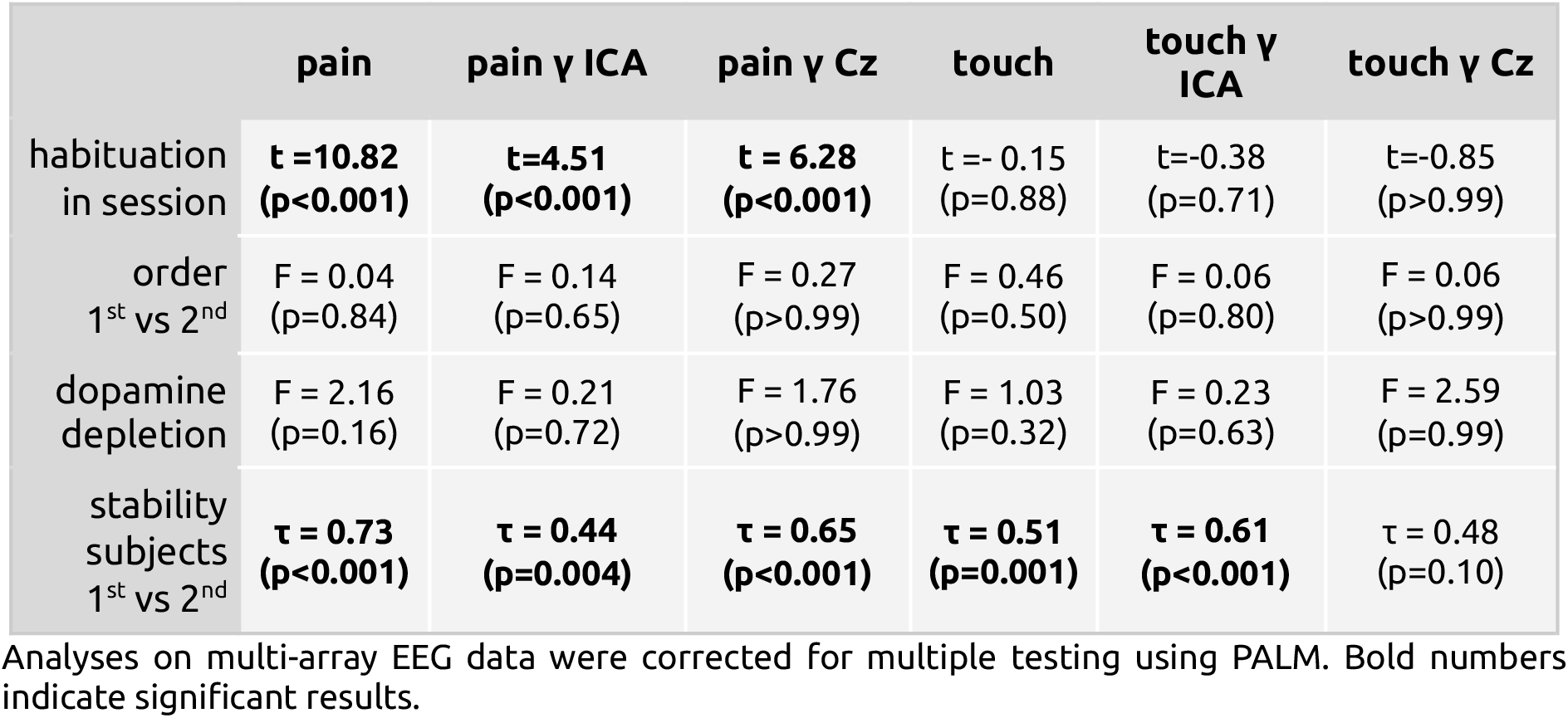
Analysis of stability, order, dopamine depletion, and habituation.

### Behavioural Data

#### Descriptive statistics

All participants received stimuli with the same physical intensity. The rating across all sessions and subjects was 4.06±1.93 for pain and 3.31±0.91 for touch (mean±std).

#### Stability

We found very consistent results of ratings between both experimental sessions. Subjects with high ratings in the first session had similarly high ratings in the second session; this applied to pain (τ = 0.73, p<0.001; Figure 2A) and touch stimulation (τ = 0.51, p=0.001; Figure 2D).

**Figure 2.**
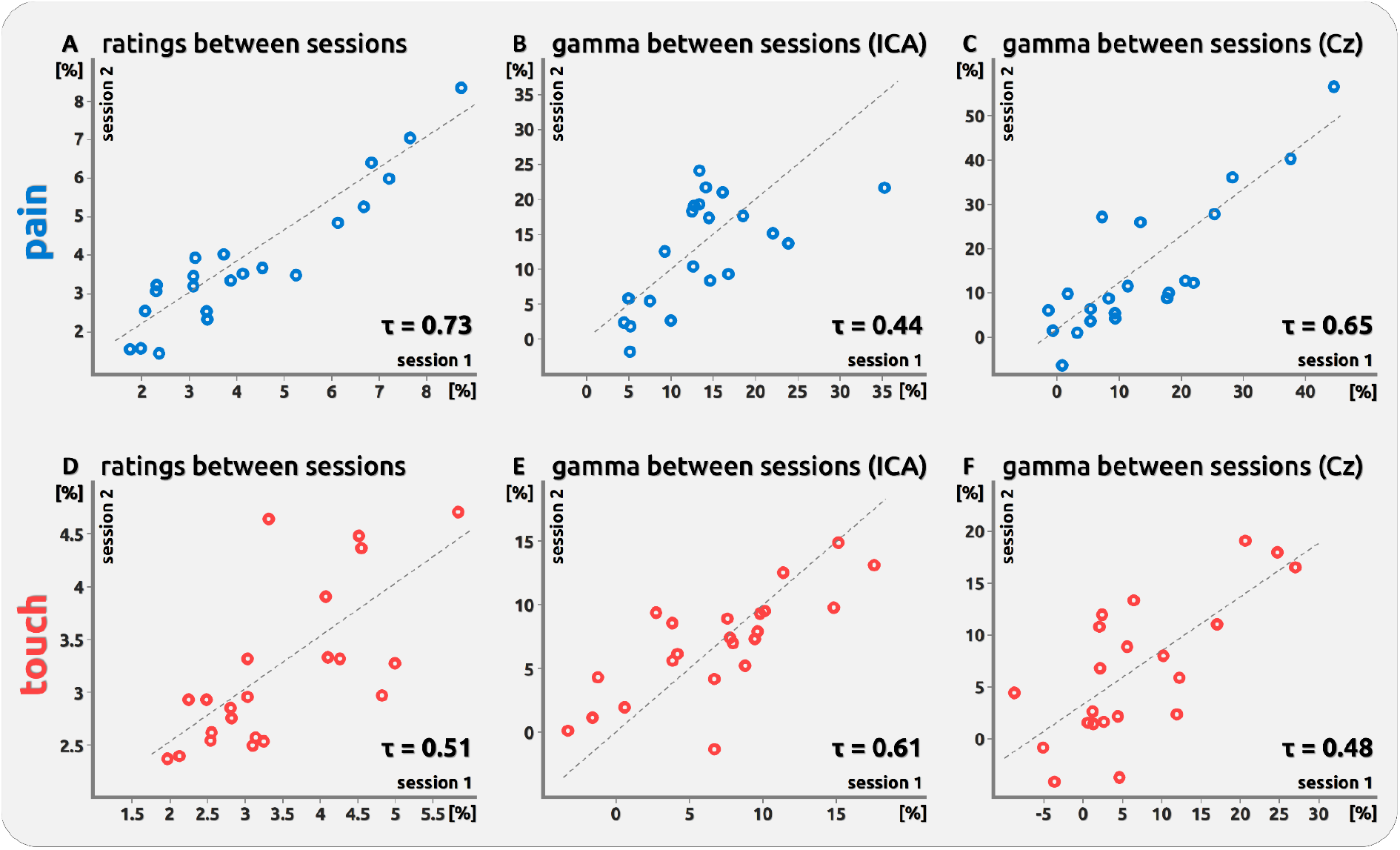
Scatterplots of individual ratings and evoked gamma. All scatterplots show stability across the two sessions. This applies to the pain domain (upper row; A,B,C), touch domain (lower row; D,E,F), as well as to averaged ratings (left;A,D), gamma derived from ICA components (middle; B,E), and gamma derived from EEG electrode Cz (right; C,F). Stability is expressed as Kendall’s τ. Data points for ICA and EEG data are shown as % signal change compared to the prestimulus baseline.

#### Order

Ratings of pain and touch were not affected by the order of the experimental sessions (pain: F[1,19]=0.04, p=0.84; touch: F[1,19]=0.46 p=0.50).

#### Dopamine

There was no effect of dopamine depletion on pain ratings (F[1,19]=2.16, p=0.16) and touch ratings (F[1,19]=1.03, p=0.32).

#### Habituation

We found an overall decrease of perception for pain trials (t = 10.82, p<0.001) but not for touch trials (t = −0.15, p=0.88). However, this effect did not apply to every participant (see distribution in Figure 3).

**Figure 3.**
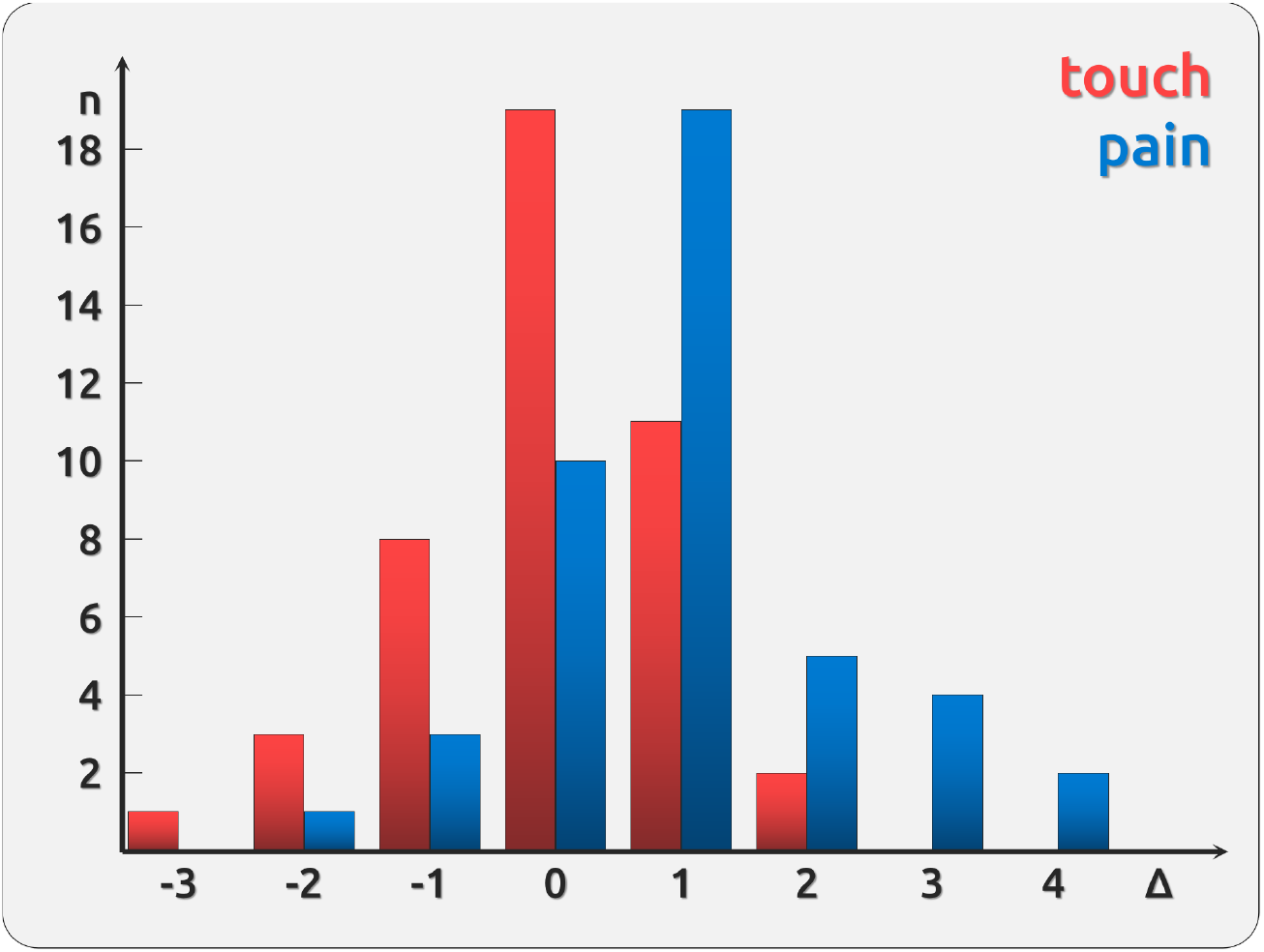
Habituation across subjects. The histogram represents the difference between the first and the last trial (Δ) based on the slope of a least-squares fitted line for each recording, e.g. 2 sessions exhibit a difference in pain ratings of about 4 between the 1^st^ and the 75^th^ pain trial; one session showed a difference of −3 between the 1^st^ and the 75^th^ touch trial. Higher differences indicate more intense first trials.

### ICA Data

#### Stability

Similar to EEG data, the magnitude of gamma response in ICA space did not differ across both experimental sessions. This applied to both pain (τ=0.44, p=0.004; Figure 2B) and touch (τ=0.61, p<0.001; Figure 2E). The TF plots and topographies can be found in Supplementary Figure 1.

In ICA space, we observed consistent TF patterns of gamma activity between both experimental sessions for the components that contain a gamma response: a “spatial” correlation of the TF pattern (range 40 to 100 Hz, 0 to 500 ms) between the first and second session showed a stronger correlation for the within-subject correlation (the diagonal in Figure 4) compared to the correlation for the subjects’ first recordings with the averaged second recordings of the other subjects (correlation coefficients outside the diagonal). This applied to gamma responses to pain (paired t-test, t=4.91, p<0.001) and touch (paired t-test, t=4.03, p<0.001). The correlation coefficients were Fisher-z transformed before the t-test.

**Figure 4.**
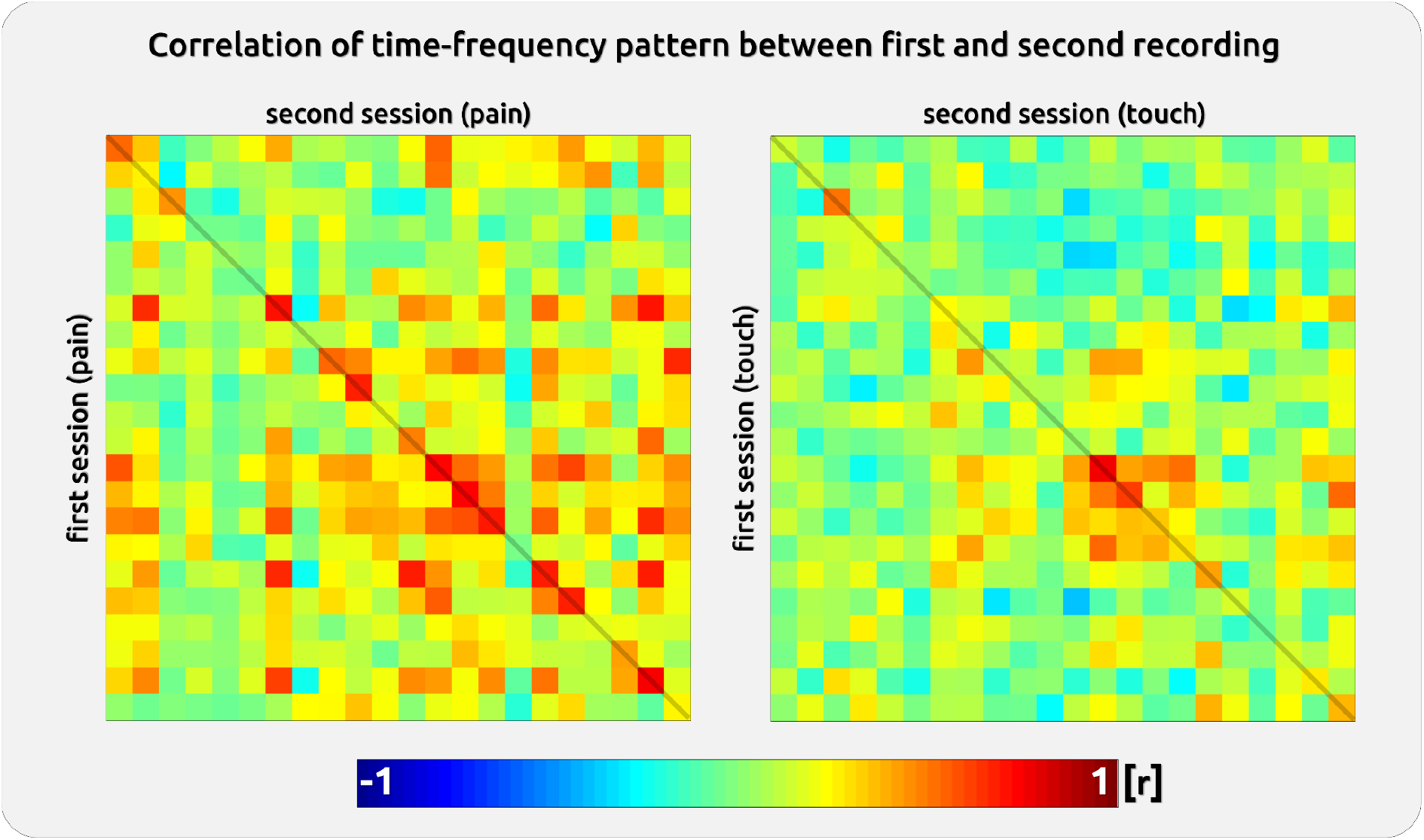
Single subject components with gamma response. Left: Correlation between the time-frequency pattern (strongest gamma component; 0-500ms / 40-100Hz) for the first session of a subject with his own and all other subjects’ time-frequency patterns Right: Same for touch stimuli. Each of the 22 rows/columns represents a single subject. The highest correlation was usually found within a subject (see diagonal). For subjects with low gamma (e.g. subject 4), the pattern correlation must inevitably fail. The correlation within a subject (diagonal) was higher than the correlation between the first session of a subject with the second session of all other subjects (everything but the diagonal). Please note that the confusion matrix is not perfectly symmetrical along the diagonal because we always correlated the first with the second recording (within and between subjects); each field is a unique correlation. Individual TF patterns are shown in Supporting Figures 1a and 1b.

#### Order and dopamine depletion

There was no significant effect for either pain or touch (Table 1).

#### Habituation

There was no significant habituation in the gamma range for touch (t=-0.38, p=0.71, df=3263) but for pain (t=4.51, p<0.001, df=3257).

### EEG Data

#### Stability

We found very consistent results of gamma activity between experimental sessions. Subjects with a high gamma in the first session had similar high gamma in the second session; this applied to pain (electrode Cz: τ=0.65, p<0.001; Figures 2C and 2), but not to touch stimulation (electrode Cz: τ=0.48, p=0.10; Figures 2F and 5). This effect was mainly distributed over the vertex and at centro-parietal electrode sites.

**Figure 5.**
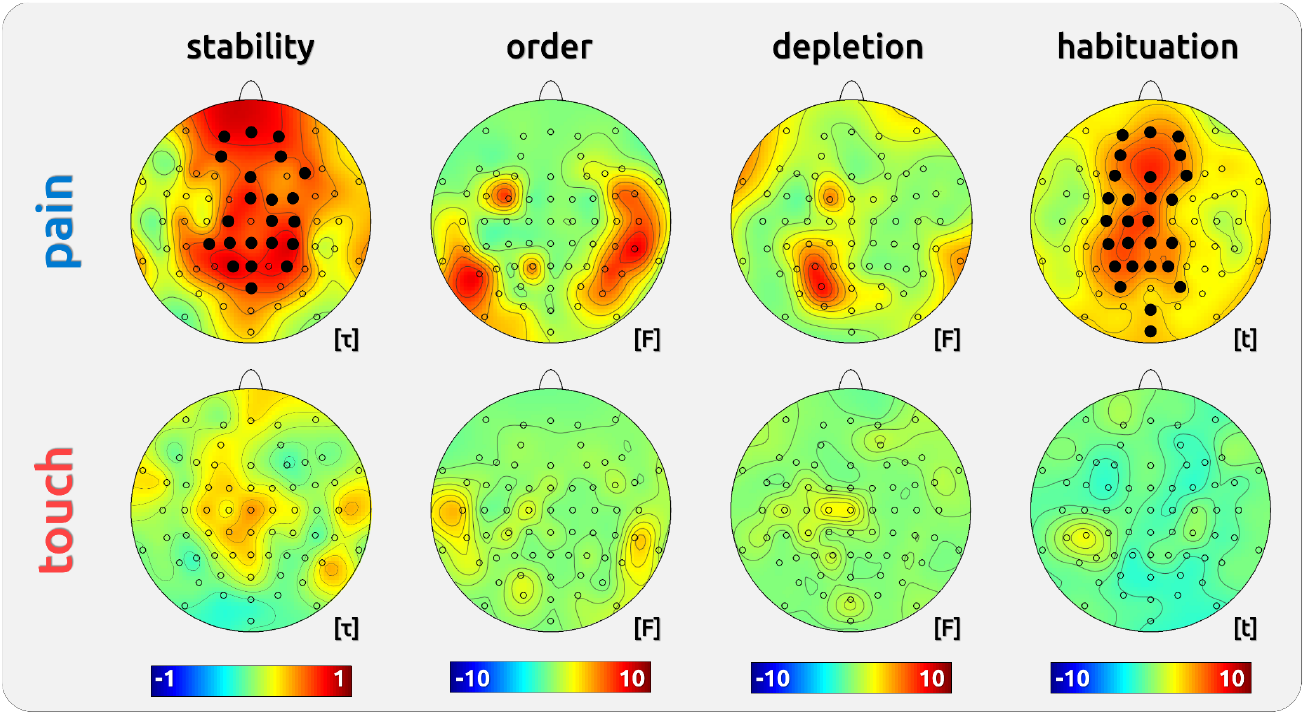
Topographies of experimental variables on gamma. Gamma responses were based on a predefined time-frequency window (150-350ms; 76-86Hz). Solid dots indicate electrodes with significant effects (p<0.05; PALM corrected). Averaged gamma responses to both stimuli and stimulus intensity encoding are shown in Supplementary Figure 1.

#### Order

There was no effect of order of stimulation sessions on gamma responses recorded at scalp vertex for both pain (electrode Cz: F[1,19]=0.27, p>0.99; Figure 5) and touch (electrode Cz: F[1,19]=0.06, p>0.99; Figure 5).

#### Dopamine

There was no effect of dopamine depletion on gamma responses for both pain (electrode Cz: F[1,19]=1.76, p>0.99; Figure 5) and touch (electrode Cz: F[1,19]=2.59, p=0.99; Figure 5).

#### Habituation

We found a drop of gamma response magnitude throughout the time course of the experiment for pain trials (electrode Cz: t=6.28, p<0.001, df=2990; Figure 5), but not for touch trials (electrode Cz: t=-0.85, p>0.99, df=2940; Figure 5). The relationship between pain intensity and gamma amplitude implies lower gamma amplitudes at the end of the experimental session.

## Discussion

Using two different levels of analysis (sensors and ICs), we aimed to investigate the individual stability of gamma oscillations time-locked to phasic noxious laser stimuli and non-noxious touch stimuli. Importantly, we delivered both pain and touch stimulation across repeated sessions within the same subject. This allowed us to unravel the relationship between perception and the magnitude of gamma responses at the individual level. Indeed, the analysis of stability compounded by the analysis of the effect of order of experimental administration, revealed that participants with high ratings and gamma amplitudes in the first session exhibited high responses in the subsequent session and vice versa (Figs. 2 and 5; Table 1). The strength of the gamma response stability at group level was maximal at the central and fronto-central region of the scalp for painful stimuli (Fig. 5, top). In addition, we found a broad variety of neuronal responses in the gamma range, ranging from clear local maxima at lateral and/or fronto-central electrodes in some subjects, to a complete absence of any gamma response in others (Supplementary Figure 1).

### Individual stability of cortical gamma oscillations

Our statistical approach allowed us to provide a robust estimate of individual gamma EEG synchronisation by integrating the information gathered by the three complementary analyses, i.e. (a) the effect of order (using LMEs), (b) between-subject signal correlation across sessions (using Kendall’s τ), and (c) the comparison of the time-frequency patterns across sessions. Indeed, whilst there was no indication of impact the order of the administration had on signal change, we found that participants displaying high gamma activity in the first session also had high gamma activity in the second session and vice versa (Figure 2).

Nevertheless, stability and reliability of EEG parameters (including gamma oscillations) have been investigated before [Fründ et al., 2007; Gasser et al., 1985; Keil et al., 2003], yet to date no study has provided evidence of an individual time-frequency estimation of acute pain-related EEG gamma responses. With a time interval of two weeks between recordings, we show remarkable stability of individual stimulus-locked high frequency EEG magnitude. Our approach may pave the way to an advanced assessment of pain-related gamma oscillations that will e.g. entail connectivity features to decode individual signature responses by means of machine learning models [Lee et al., 2021; Schulz et al., 2012].

Importantly, cortical patterns associated with brief touch and pain stimulation deviated substantially from the gamma topography of the group statistics. Hence group analyses involving a single electrode would likely underestimate the distribution of the experimental effects as some participants will display local maxima at other electrodes. In a similar vein, the results of group statistics implicitly suggest that gamma oscillations reflect pain perception in every participant [May et al., 2018; Schulz et al., 2011; Zhang et al., 2012], which is not the case. In fact, we show how participants exhibited more than one gamma response pattern, characterised by non-overlapping time-frequency activity and distinct topographies, especially in the pain condition. Previous findings suggesting that gamma brain oscillations mediate perception of pain would benefit from a repeated recording in order to confirm the stability of individual activity patterns [Liberati et al., 2018].

### Analyses on reproducibility may neglect the individual differences

The present data suggest that group results on gamma oscillations are replicable. Gamma responses to laser pain have been shown in a number of studies [Bassez et al., 2020; Hauck et al., 2015], however, a simple repetition of findings at group level can be misleading. First, group studies do not sufficiently take into account the variability of the individual. For example, the interpretation of group analysis implies that every participant shows the gamma effect at 80Hz, but the time-frequency variability of the gamma response and its spatial distribution is such that group analyses eventually fail in accounting for different individual patterns (such as the absence of gamma). Studies may also overestimate the contribution of a cortical process to pain perception. The present study confirms further evidence that the results of group statistics do not necessarily reflect the cortical processing of the individual [Mayr et al., 2020; Mayr et al., 2021]. This finding has implications for studies implementing haemodynamic measures. For example, neuroimaging studies reporting the posterior insular activity as a crucial node of the network of brain structures generating the experience of pain may fail to quantify individuals with very low or no activation of this structure. Future studies with multiple repeated recordings of a single participant are needed to shed light on this widely neglected issue.

### Conclusions

The present study aimed to investigate the stability of pain-related gamma oscillations across repeated recording sessions. To the best of the author’s knowledge, this is the first study to show remarkable stability of the phasic pain-related time-frequency gamma oscillations for the individual. These individually stable patterns are not reflected by group statistics and would require a more thorough investigation. In fact, the importance of assessing stability is equally reflected by the large variability of pain-related gamma oscillations. Future research may aim to elucidate the conditions under which some subjects do not exhibit pain-related gamma oscillations at all or whether some individuals would display stable cortical processes for long-lasting tonic and chronic pain.

In conclusion, the present results support the notion of co-occurring intra-individual stability and inter-individual variability of neural gamma synchronisation and argue fora greater emphasis on subject-level assessment of sensory responses, particularly in the context of pain perception.

## Acknowledgements

We thank Dr Stephanie Irving for copy-editing the manuscript.

## Supporting Material

**Supporting Figure 1a.**
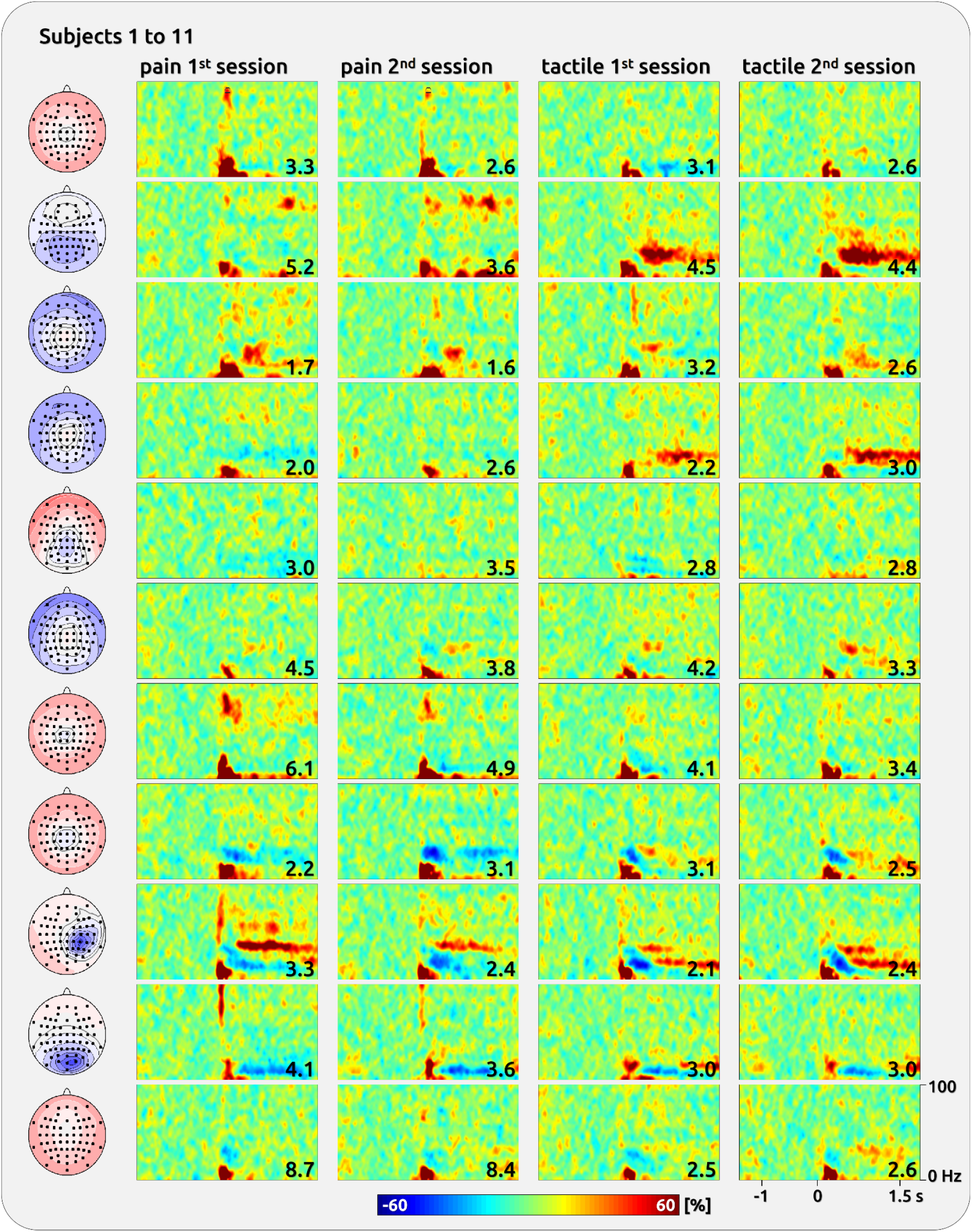
Single subject components with gamma response. Left: topographies of ICA components. Right: TFR separately for the first and second session and for pain and touch. The numbers represent the averaged ratings.

**Supplementary Figure 1b.**
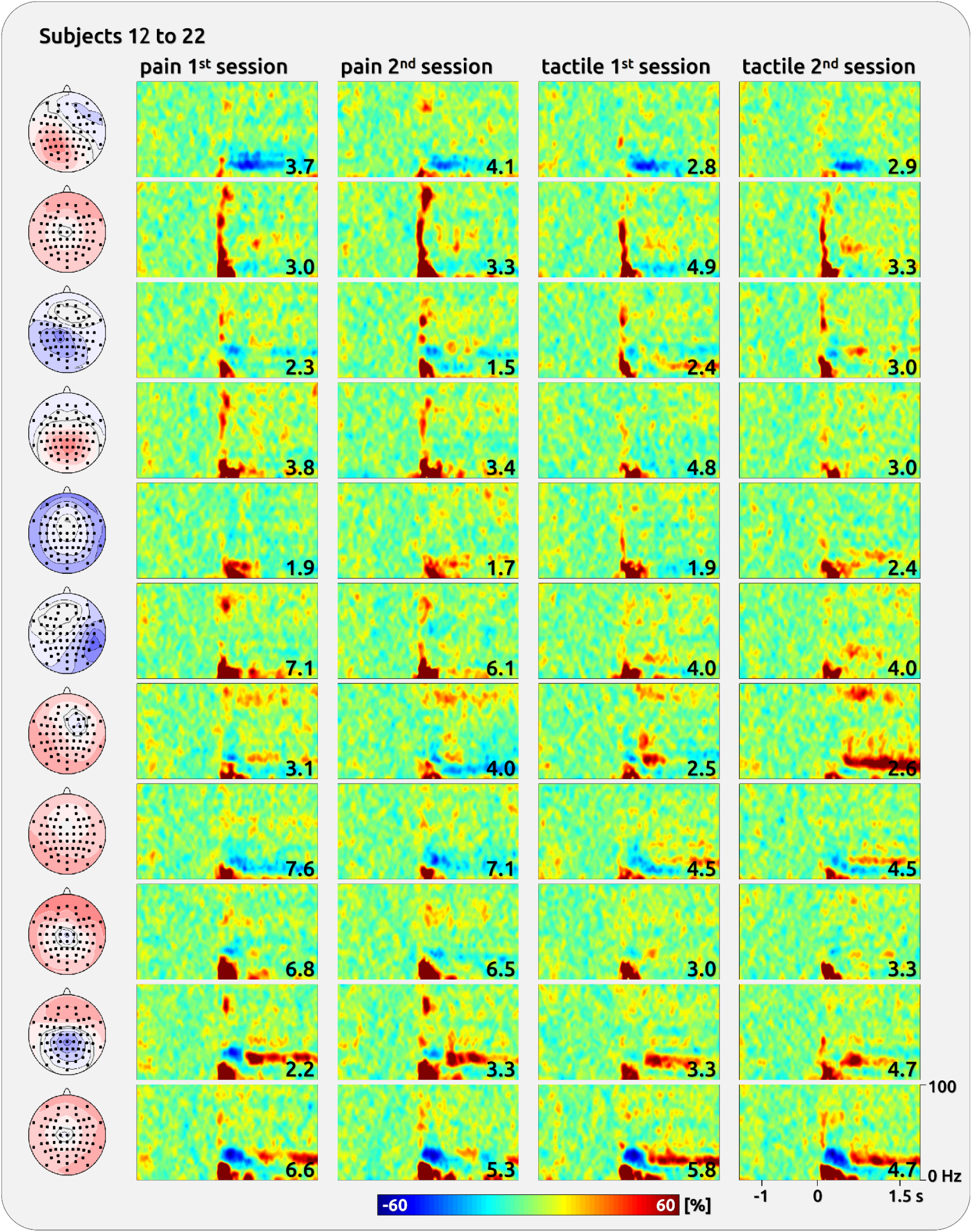
Single subject components with gamma response. Left: topographies of ICA components. Right: TFR separately for the first and second session and for pain and touch. The numbers represent the averaged ratings. The topographies were normalised and range from −1 to 1.

