## Supporting Material for "Individual Stability of Pain- and Touch-Related Neuronal Gamma Oscillations"

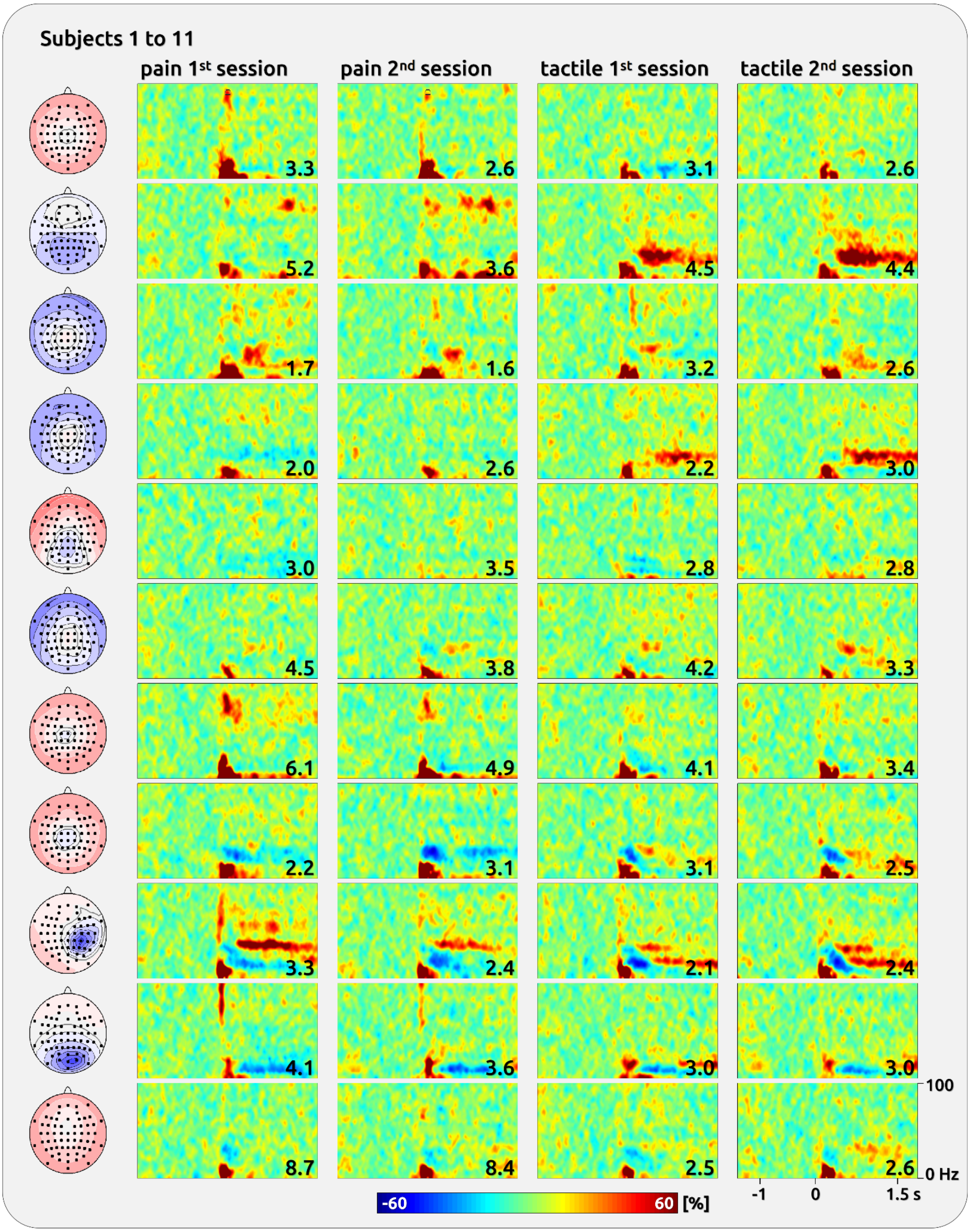


**Supporting Figure 1a | Single subject components with gamma response.** Left: topographies of ICA components. Right: TFR separately for the first and second session and for pain and touch. The numbers represent the averaged ratings.

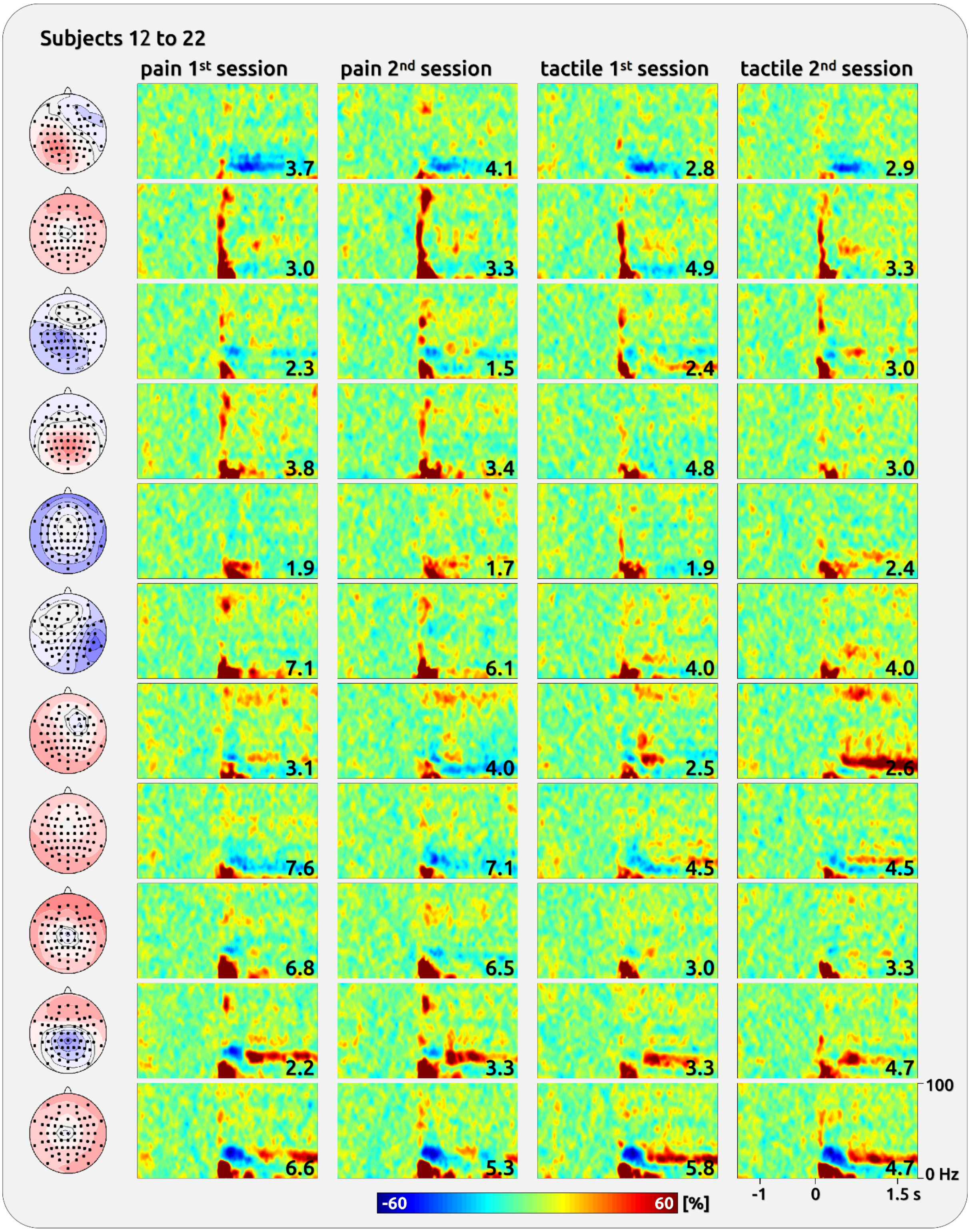


**Supplementary Figure 1b | Single subject components with gamma response** Left: topographies of ICA components. Right: TFR separately for the first and second session and for pain and touch. The numbers represent the averaged ratings. The topographies were normalised and range from -1 to 1.
